# From Diversity to Dominance: How Salt and CO_2_ Shape LAB-dominated Ecosystems in Vegetable Fermentations

**DOI:** 10.1101/2025.10.11.681785

**Authors:** Tom Eilers, Tim Van Rillaer, Stijn Wittouck, Ines Tuyaerts, Katrien Michiels, Maline Victor, Thies Gehrmann, Peter A Bron, Wannes Van Beeck, Sarah Lebeer

**Affiliations:** Department of Bioscience Engineering, Lab of Applied Microbiology and Biotechnology, University of Antwerp, Groenenborgerlaan 171, 2020 Antwerp, Belgium

## Abstract

Research on microbial ecosystems is often challenging due to high diversity of microbial taxa present and the complexity of controlling environmental variables. To address these challenges, fermented foods are simpler and more reproducible model ecosystems, where both community composition and environmental factors can be more precisely controlled and manipulated. In this study, we focused on fermented vegetables which are typically dominated by lactic acid bacteria (LAB). It is not completely understood why lactic acid bacteria (LAB) consistently drive the spontaneous fermentation of vegetables such as cabbage and carrots and how variables such as vegetable substrates, salt addition, and carbon dioxide levels can impact microbial community dynamics. Here, we explored the temporal microbial dynamics in standardized fermentations of 11 different vegetables (including beetroot, bell pepper, cabbage, carrot, cucumber, fennel, green asparagus, leek, parsnip, sunroot, and tomato), revealing a consistent dominance of *Leuconostoc* and other LAB. Additionally, we investigated the impact of varying salt concentrations, demonstrating that lower salt levels resulted in a delayed appearance of the typically dominant LAB community, while simultaneously revealing a higher abundance of *Weissella* and various *Enterobacterales* taxa. These effects imposed by reduced salt concentrations were mitigated by CO_2_ injection, which reverted the enhanced *Enterobacterales* levels and increased the overall abundance of *Lactobacillales*. This study demonstrates how targeted manipulation of environmental parameters, such as salinity and gas composition, can be used to uncover ecological principles governing microbial succession and community assembly in reproducible fermentation-based model ecosystems.

**Importance:** Understanding the ecological principles that shape microbial community assembly is essential for advancing our knowledge of microbial ecosystems. Fermented vegetables, increasingly popular among the general population, provide tractable and reproducible model systems to study microbial succession under controlled environmental conditions. By systematically manipulating variables such as vegetable type, salinity and gas composition, we uncovered the effect of these factors on the microbial dynamics throughout the fermentation. These insights not only contribute to a better understanding of the microbial ecology of these man-made food systems but also suggest directions for novel strategies to optimize fermentation processes for producing faster, safer, and more flavorful foods.

## Introduction

Research on microbial ecosystems is often challenging due to the large diversity of taxa and the numerous and often unknown external factors that shape ecosystem development. To better understand the key drivers of microbial dynamics and interactions, simpler, manipulable and reproducible model systems are needed. Fermented foods offer such systems, natural yet tractable ecosystems, in which external parameters can be systematically varied^1^. Their adaptability allows researchers to modulate conditions such as substrate type, salinity, CO_2_ levels, temperature, and fermentation duration. This type of mechanistic ecological research not only advances our understanding of microbial community assembly but also informs practical strategies for microbiome engineering in applied contexts.

In recent years, fermented food ecosystems have also gained renewed popularity in society, driven by the growing production and consumption of artisanal and industrial fermented products. This resurgence in Western culture is fueled by their desirable organoleptic properties, their potential to promote sustainability by reducing food waste, and increasing evidence of their health benefits ^2–6^. Fermented foods encompass distinct food products, but here we adhere to the scientific definition that fermented foods are ‘foods made through desired microbial growth and enzymatic conversions of food components’ ^7^. It is estimated that over 5000 different fermented foods exist globally, which can be divided based on the substrate fermented ^8^. In addition to cow’s milk, vegetables form one of the most important groups of substrates of fermented foods ^9^.

Vegetable fermentation typically begins with chopping or crushing the vegetables, followed by the addition of salt and the exclusion of air by sealing the mixture in airtight jars. These conditions selectively stimulate the growth of a desired subset of the native microbial community, primarily lactic acid bacteria (LAB) ^10^. LAB are known for their ability to ferment plant-derived sugars into lactic acid, which lowers pH and inhibits the growth of undesirable microbes, particularly members of the order Enterobacterales^11^. These spoilage-associated taxa typically co-occur with LAB at the onset of fermentation and are linked to the production of off-flavors and potentially harmful compounds such as biogenic amines and toxins^12^.

A few selected vegetable fermentations have been extensively studied so far and include sauerkraut ^13,14^, kimchi ^15,16^, cucumber ^17^, and fermented carrot juice ^12,18^. Among these fermentations, cabbage and carrot fermentations consistently exhibit a characteristic succession: an initial dominance of heterofermentative LAB, such as *Leuconostoc*, followed by homofermentative genera like *Lactiplantibacillus* and *Latilactobacillus*^12,13,19^. However, it remains unclear whether similar microbial dynamics occur in the fermentation of other vegetables. Moreover, the ecological mechanisms driving LAB dominance and succession are still poorly understood. Potential drivers include the starting substrate’s initial microbial community ^20^ and nutrient composition, ^21^, as well as broader environmental influences, akin the concept of ‘microbial terroir’ in wine fermentation, where geographic and environmental factors shape microbial communities ^22^. In this context, it is still unknown whether the location on the plant, i.e. soil-exposed, nutrient-dense rhizosphere and air exposed phyllosphere and carposphere affect the microbial dynamics during fermentation.

Other environmental variables affect and can thus be modulated to influence the microbial community dynamics that occur during the fermentation. Traditionally, variable concentrations of salt, typically ranging from 2-8%, are added to vegetable fermentations ^23–25^, decreasing the water activity ^26^, increasing osmotic stress. However, such high salt concentrations are negatively perceived by consumers for organoleptic properties ^27^, and linked to (presumed) negative health effects of the fermented end product ^26,28,29^. Recent studies have explored reducing salt concentrations in traditional fermentations, down to 1.4% in kimchi ^19^, 0.5% in sauerkraut ^30^, and 0% in cucumbers ^31^, while monitoring metabolites and pH dynamics. Although these studies only partially examined the microbial communities, they reported distinct shifts in microbial composition and fermentation profiles. However, comprehensive investigations into the complete removal of salt and its effects on the full microbial community remain scarce.

Atmospheric conditions, including the presence of carbon dioxide (CO_2_), represent another environmental factor influencing microbial community dynamics during fermentation. Carbon dioxide (CO_2_) has been shown to inhibit a broad range of spoilage organisms. For instance, in liquid broth experiments where CO_2_ was introduced via gas packaging, concentrations of 2000 mg/L CO_2_ completely inhibited several Gram-negative bacteria such as *Pseudomonas fluorescens, Photobacterium phosphoreum, Shewanella putrefaciens*, and *Aeromonas hydrophila*. Gram-positive such as *Brochothrix thermosphacta* and *Bacillus circulans* ^32^ were also inhibited, albeit to a lesser extent at the same concentration. In contrast, the LAB *L. sakei* exhibited only a moderate reduction in growth rate at concentrations up to 2500 mg/L CO_2_, suggesting that CO_2_ may act as a selective pressure favoring LAB dominance. Interestingly, CO_2_ is also naturally produced during vegetable fermentation by heterofermentative LAB such as *Leuconostoc* spp., yet its potential role in shaping microbial succession remains unexplored. Understanding how atmospheric composition influences microbial ecology could open new avenues for steering fermentation dynamics, particularly in low- or no-salt conditions.

To clarify how substrate characteristics, salinity, and atmospheric conditions (in particular CO_2_) shape microbial community assembly and succession in plant-associated fermentation ecosystems, we conducted a systematic investigation under standardized conditions. Eleven vegetables (Table 1) coming from distinct plant regions, carposphere (e.g. tomato), phyllosphere (e.g. cabbage), and rhizosphere(e.g. carrots and sunroot), were fermented, resulting in 228 samples collected at two timepoints. The active microbial communities were analyzed at the genus level using the updated *Lactobacillaceae* taxonomy ^33^. The specific effects of salt and CO_2_ were examined using carrot juice fermentations as a controlled model system, enabling detailed analysis of their impact on microbial succession and LAB dominance.

**Table 1.** Overview of the various vegetables used in this project.

| Vegetable | Latin name | Plant organ | Region of plant |
| --- | --- | --- | --- |
| Beetroot | *Beta vulgaris* var. Ruba | Taproot | Rhizosphere |
| Carrot | *Daucus carota* subsp. *sativus* | Taproot | Rhizosphere |
| Parsnip | *Pastinaca sativa* | Taproot | Rhizosphere |
| Sunroot | *Helianthus tuberosus* | Tuber | Rhizosphere |
| Bell pepper | *Capsicum annuum* | Fruit | Carposphere |
| Cucumber | *Cucumis sativus* | Fruit | Carposphere |
| Tomato | *Solanum lycopersicum* | Fruit | Carposphere |
| Cabbage | *Brassica oleracea* var. *capitata* | Leaves | Phyllosphere |
| Leek | *Allium ampeloprasum* var. *porrum* | Stem and leaves | Phyllosphere |
| Fennel | *Foeniculum vulgare* | Stem and leaves | Phyllosphere |
| Green asparagus | *Asparagus officinalis* subsp. *officinalis* | Modified stem | Phyllosphere |

## Materials and Methods

### Spontaneous vegetable fermentations and sampling

Vegetables (Table 1) were sliced and fermented in 2.5% w/v NaCl solution, following a previously established protocol for carrot fermentation^12^. Fermentations were conducted under controlled conditions, within a temperature-controlled room at 20°C. 400 µl of sample were taken on day 7 and 14, and frozen in liquid nitrogen and stored at -80°C for subsequent RNA extraction to measure the active community.

### Varying salt and CO_2−_levels in lab-controlled vegetable fermentations

To determine the influence of addition of salt and CO_2_ on the progression of vegetable fermentations, carrot juice fermentation was monitored over a 30-day period. Carrots were rinsed with H_2_O and were processed with Solis Juice Fountain Pro Type 843 centrifuge. Varying salt concentrations were added to the fermentation of 0%, 1.25% or 2.5% w/v NaCl. Carbon dioxide (CO_2_) was introduced by saturating the samples using a standard home kitchen SodaStream (SodaStream), and the fermentations were divided into 0.25 l Weck jars. To minimize interference, individual temporal samples were collected from separate Weck jars, at timepoints 0, 2, 3, 7, 14, and 28 days. Each 400 µl sample was immediately frozen in liquid nitrogen and kept at -80°C for subsequent RNA extraction.

### V4-16S rRNA amplicon sequencing of the active microbial fraction

Total RNA was extracted using the RNeasy PowerMicrobiome kit (Qiagen), which included an on-column DNase digestion step. After RNA extraction, the absence of DNA was checked using a PCR on the 16S region (with 27F and 1492R), and the lack of PCR product was confirmed on a 1% agarose gel. Complementary DNA (cDNA) was synthesized using Readyscript (Sigma-Aldrich), and 16S rRNA amplicon sequencing was performed as previously described in De Boeck et al. 2020 ^12^.

### Data analysis

The sequencing results were processed with the DADA2 package, version 1.22.0 ^34^, as described in Lebeer et al (2023). However, reads were classified up to genus level using a combined database from GTDB (version r207 ^35^) and the SILVA ribosomal RNA gene database (SILVA SSU 138.1 ^36^), which already included the novel *Lactobacillales* taxonomy as described by Zheng et al (2020) ^33^. Notably, a suffix behind the genus name, for instance Lactococcus_A, indicates that according to the GTDB the taxon it was named after (without the suffix) was too diverse and/or not monophyletic. In this case, Lactococcus_A refers to the *Lactococcus carnosus* clade.

Subsequent data analysis and interpretation were executed in the R environment using in-house developed packages including tidytacos (github.com/LebeerLab/tidytacos, “tidy Taxonomic Compositions”) ^37^, which builds on tidyverse ^38^ specifically for the exploration of microbial community data. In addition, it includes multidiffabundance ^39^ (github.com/thiesgehrmann/multidiffabundance/) ^39^, which builds on various tools for microbiome analysis, including differential abundance tools ALDEx2^40^, ANCOM-BC2^41^, corncob ^42^, DESeq2^43^, limma^44^, lmclr ^39^, maaslin2^45^, and ZicoSeq ^46^. When multiple testing was performed, p-values were adjusted for false discovery rates.

## Results

### *Leuconostoc*-dominance is conserved across a wide variety of vegetable fermentations

Microbial fermentations offer a simplified yet dynamic system to study ecological assembly processes. In this study, we examined how different vegetable substrates shaped microbial community structure and succession by fermenting 11 commonly consumed vegetables (Table 1). 114 vegetable fermentations were set up in a standardized way, with addition of 2% NaCl and sampled after 7 and 14 days for microbial community analysis (Figure 1A). At the end of these 14day-fermentations, asparagus showed the highest alpha diversity (inverse Simpson: 4.92 ± 0.76, Figure 1C) among all fermented vegetables tested, whereas sauerkraut and fennel showed the lowest alpha diversity (inverse Simpson: 1.33 ± 0.45 and 1.86 ± 0.37, respectively).

**Figure 1.**
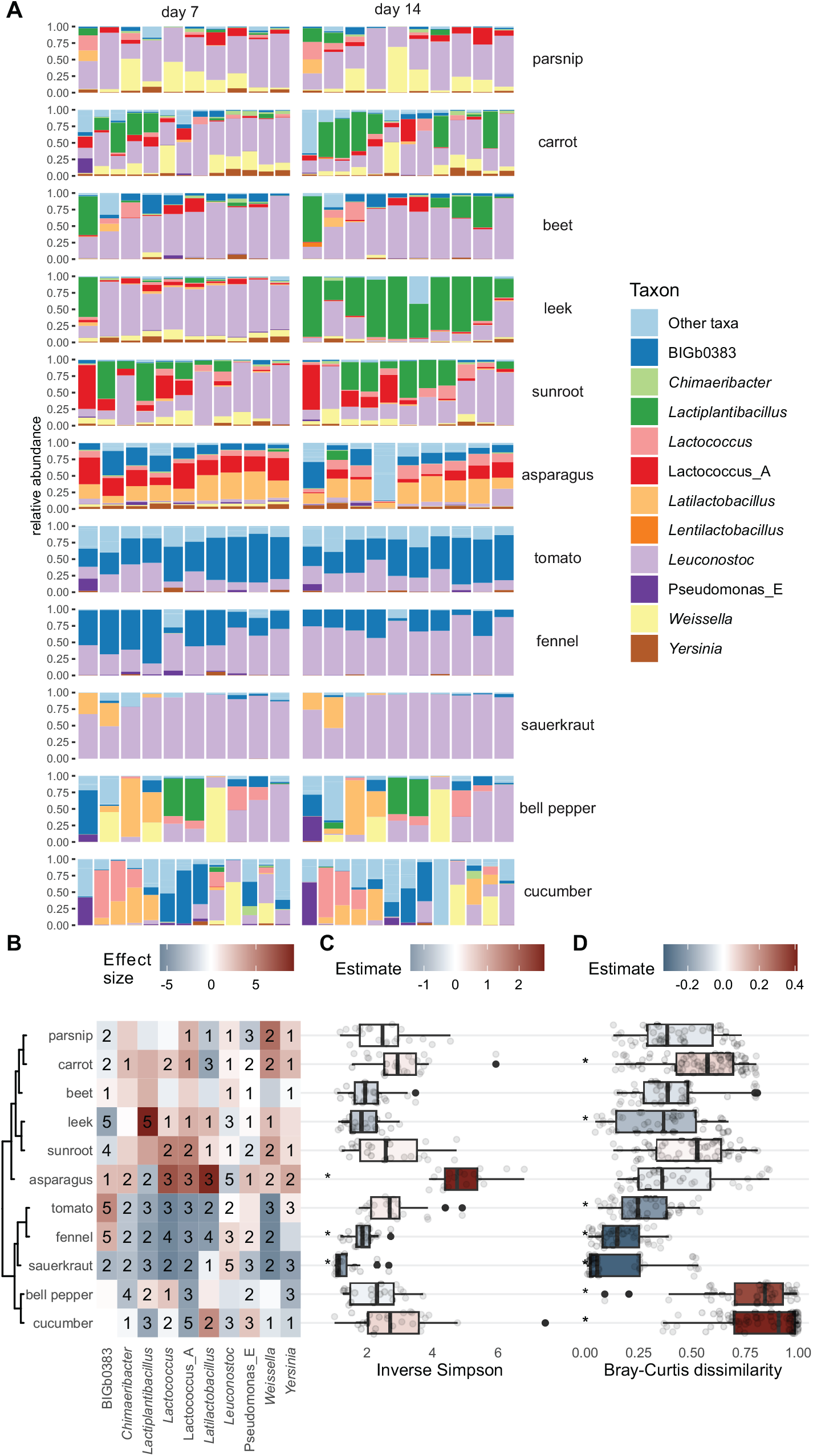
Active bacterial community structure of different vegetables fermented with 2 % salt added. Vegetable fermentations were ordered based on a hclust on the Maaslin2 effect sizes. A: Overview of the relative abundances of the different vegetable fermentations on day 7 and day 14. The bar plot illustrates genus relative abundances in samples from day 7 and 14. B: Differential abundances of the top 11 most dominant taxa across all fermentations assessed with 8 different differential abundance metrics for each vegetable tested against the collective group. Color shows maaslin2 effect size. Number represent the total number of statistical tests that were significant with FDR corrected p-value <0.05. C: Alpha diversity for each vegetable. D: pairwise Bray-Curtis dissimilarity within vegetables. The color shows the direction of the effect size compared to all other vegetables, p-value <0.05 “*”.

LAB appeared highly prevalent and abundant in most of the fermentations studied, with *Leuconostoc* the most abundant genus found in 117 of 228 fermentation samples analyzed. Other prominent gena included *Lactiplantibacillus* (29/228), *Latilactobacillus* (18/228), and *Weissella* (10/228). Notably, an uncharacterized *Enterobacteriaceae* taxa BIGB0383 belonging to the family *Enterobacteriaceae* (Supplementary S1), was the most abundant genus in 35 of the 232 fermentation samples and significantly more abundant in tomato and fennel. Differences in microbial diversity and composition were also observed between the different fermented vegetable substrates studied. An adonis test demonstrated that the type of vegetable substrate used significantly influenced overall bacterial composition across all vegetable fermentations. Specifically, the region of plant (Carposphere, phyllosphere or rhizosphere) explained 6.61% of the variance on day 7 and 9.56% on day 14. More importantly, the specific vegetable fermented could explain 22.06% on day 7 and 22.68% on day 14. Since the overall community composition was not significantly different between time points across all vegetables, only data from day 14 were included in the remainder of the analyses (R^2^ = 0.0017, p=0.8).

To more specifically explore the differences in microbial community between the different vegetables, we performed a differential abundance analysis using eight statistical methods (Figure 1B). While most fermentations were dominated by LAB, the most predominant distinction observed on day 14 was that fermented leek showed a higher abundance of *Lactiplantibacillus* and lower abundance of BIGB0383 compared to the other vegetables. In contrast, BIGB0383 was more abundant in all tomato and fennel fermentations and was also present in some cucumber and asparagus fermentations, albeit at lower abundances (Figure 1B). Fermented green asparagus and tomato had significantly higher abundances of *Yersinia*, while sauerkraut and bell pepper had a lower abundance of this potential pathogen. Remarkably, fermented green asparagus showed a significantly higher abundance of *Latilactobacillus, Lactococcus*, and lower *Lactiplantibacillus* compared to all other vegetables (Figure 1B). Within-vegetable variation was assessed by calculating pairwise Bray-Curtis dissimilarity values (Figure 1D). These analyses confirmed that cucumber varied most across samples, while sauerkraut fermentations appeared most similar.

Despite these differential abundance profiles and differences in overall composition, a consistent pattern observed across most fermentations was that LAB, and especially *Leuconostoc*, were dominant across most fermentations, with its the composition and diversity varying depending on the vegetable substrate tested.

### Salt reduction slows down LAB dominance and increases *Enterobacterales* abundance

While the vegetable substrate plays a key role in shaping microbial communities, environmental factors must also be considered in these man-made ecosystems. Among these, salt addition is one of the primary interventions used to steer microbial dynamics in fermented vegetables. To better understand its influence, we investigated how different salt concentrations influence active microbial community composition and the potential suppression of spoilage organisms and pathogens. We selected carrot juice fermentation as a model system, based on our prior experience with this ecosystem and its practical advantages in the laboratory. Its liquid nature, in particular, makes it easier to handle and sample under controlled conditions ^12,24^. Carrot juice fermentations were set up with three salt concentrations, 2.5%, 1.25%, and 0% NaCl, to mimic standard conditions, lower salt concentration, and no salt conditions, respectively, Samples were collected over a 28-day period. As observed in the various vegetable fermentations, LAB were the most predominant members with low variation between different biological replicates. The microbial community initially consisted primarily of heterofermentative *Leuconostoc* and gradually shifted to a more homofermentative *Lactiplantibacillus-*dominated community, depending on the salt concentration (Figure 2A).

**Figure 2.**
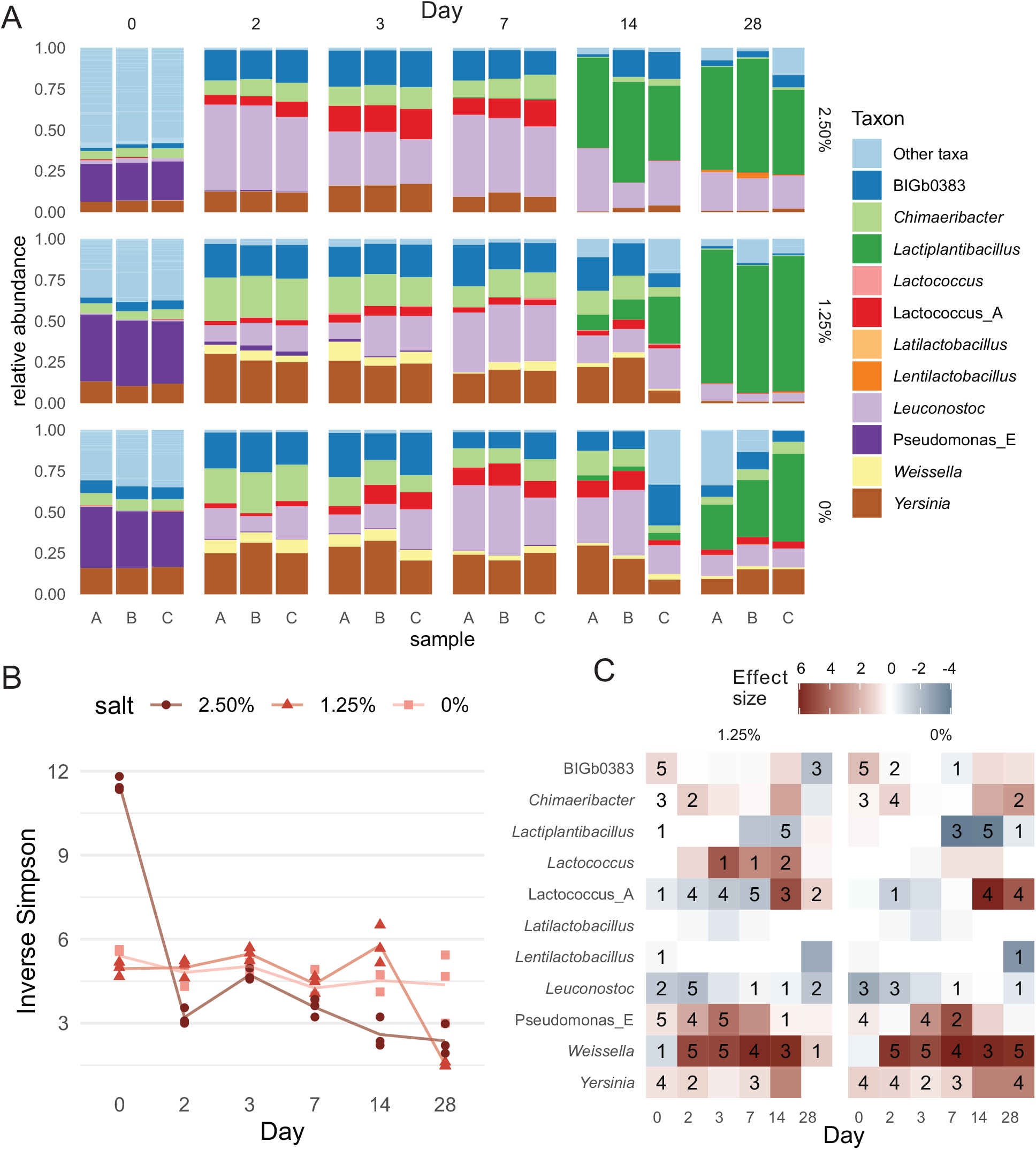
Impact of salt concentration on carrot juice fermentation. **A**: Bar plot illustrating carrot juice fermentations with three replicates of three different salt conditions over time **B**: alpha diversity calculated with inverse Simpson over time compared to the samples with a reference salt concentration of 2.50%NaCl. Different shades of red represent the different salt (NaCl) concentrations used. **C**: A heatmap comparison of the differential abundances compared to the reference salt concentration of 2.50%NaCl of the 11 most abundant taxa over time, calculated with 8 different differential abundance tests, with effect size visualized by maaslin2. (p-values were corrected for false discovery rate FDR, “*”= p <0.05). Number depicts the number of significant differential abundance tests.

Alpha diversity was used as a proxy for fermentation consistency, with lower diversity indicating a more uniform LAB-dominated community (Figure 2B). Compared to 2.5% NaCl, significantly higher alpha diversity was observed up to day 14 in fermentations with 0% and 1.25% salt. By day 28, diversity in the 1.25% and 2.5% salt groups converged, suggesting that 1.25% NaCl fermentations progressed more slowly but eventually reached a similar *Lactiplantibacillus*-dominated state. In contrast, diversity remained higher in the no-salt group, representing lower LAB dominance.

Differential abundance analysis revealed that on day 0, fermentations with all salt concentrations displayed high levels of *Pseudomonas, Yersinia, Chimaeribacter*, and the unclassified *Enterobacterales* taxon BIGb0383, though at varying abundances. When 2.5% salt was added, *Leuconostoc* became dominant (>50% relative abundance) after 2 days, while under 1.25% and 0% salt conditions, this shift occurred after 7 days (Figure 2C). A similar delay was observed for *Lactiplantibacillus*, which dominated by day 14 in 2.5% salt, but only by day 28 in lower salt conditions. Notably, in the no-salt group, only 2 out of 3 fermentations still reached *Lactiplantibacillus* dominance by day 28. Low and no-salt fermentations also showed prolonged presence of potential spoilage taxa such as *Yersinia, Pseudomonas_E, Chimaeribacter*, and *BIGb0383. Weissella* was more abundant at nearly all timepoints in low-salt conditions, while *Lactococcus_A* increased later in fermentation. Taken together, these results indicate that reducing salt hampers LAB succession and allows spoilage-associated taxa to persist longer.

### CO_2_ addition under low salt conditions restores rapid LAB community domination

When lower salt concentrations are desired, alternative strategies are required to ensure LAB dominance. Therefore, we next investigated the potential of CO_2_ as a selective intervention to support LAB-driven fermentation. Hereto, carrot juice fermentations were performed with or without prior CO_2_ injection. Notably, CO_2_-treated fermentations started a lower pH (6.08 ± 0.03 vs 5.43 ± 0.03; p < 0.05; Figure 3A), suggesting that CO_2_ injection led to an initially more acidic environment. Despite this initial decrease, the overall pH trajectory during the fermentation was not altered in later stages. Interestingly, by day 2, pH values in CO_2_-treated fermentations were slightly higher than in non-treated controls (p < 0.05), suggesting a small temporary buffering effect despite the initially lower pH. CO_2_ injection significantly influenced early fermentation dynamics. Similar to the fermentations without CO2-injection, the communities were dominated by *Leuconostoc*, followed by the homofermentative *Lactiplantibacillus*. However, in the later stages, some samples with lower salt concentration contained *Lentilactobacillus* (Figure 3B). Alpha diversity remained significantly lower in CO_2_-treated fermentations compared to controls until day 7, after which no differences were observed (Figure 3C).

**Figure 3.**
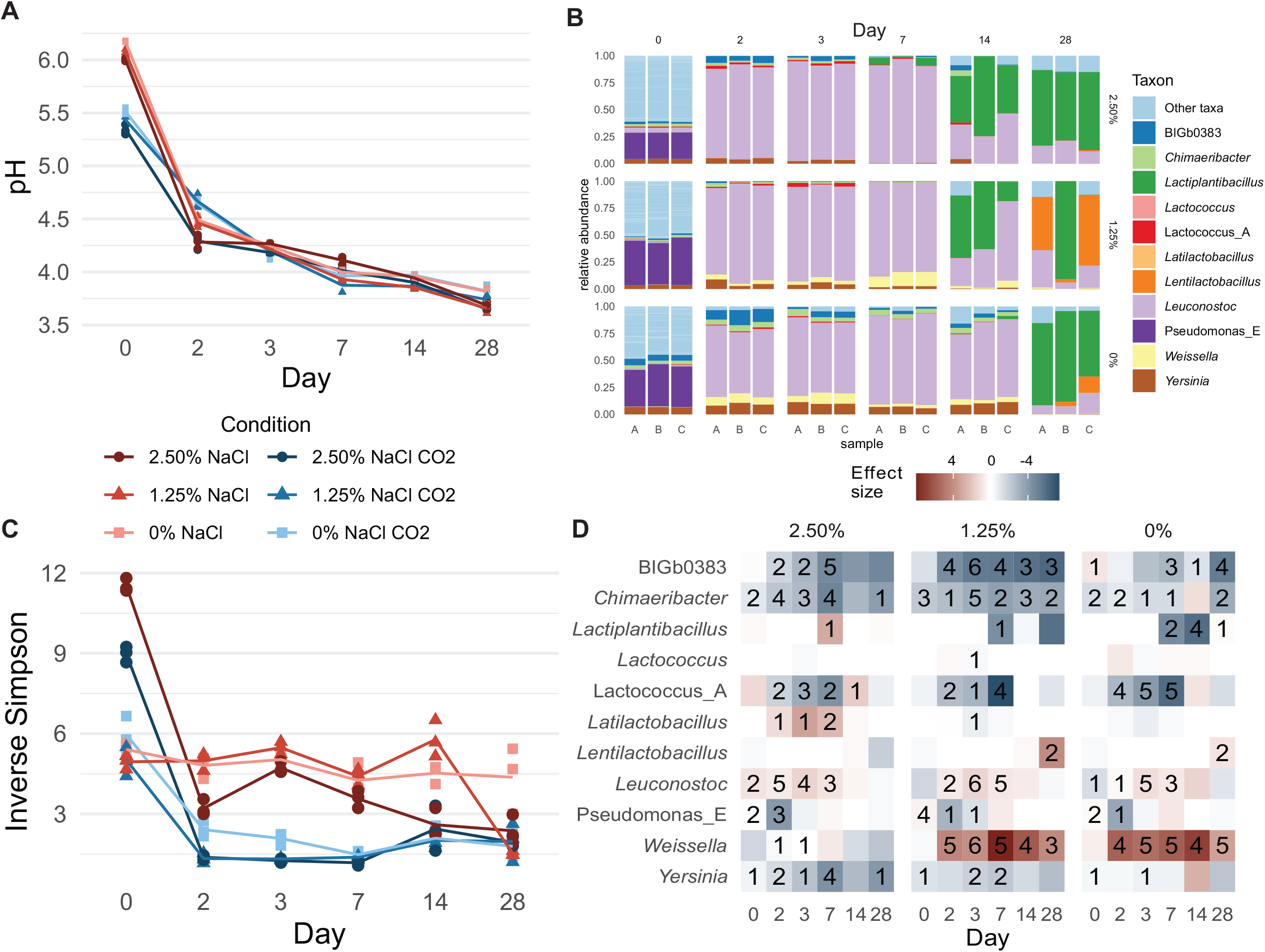
Influence of CO_2_ on microbial community dynamics in carrot juice fermentations. Shades of red present the fermentations with CO_2_ treatment (CO_2_), while shades of blue represent the fermentations without CO2 injected. **A**: pH of carrot juice fermentations with various salt concentration and addition of CO_2_ before the start of fermentation. **B**: Bar plot illustrating the microbial community after addition of CO_2_ in the fermentation under different salt concentrations, for the control, see Figure 2A. **C**: Alpha diversity over time compared to the reference salt concentration of 2.50% NaCl. **D**: Differential abundance CO_2_-injected fermentations compared to 2.5% NaCl without CO_2_ determined with Maaslin2. (“*”=FDR corrected p<0.05).

Differential abundance analysis at the genus level showed that CO_2_ injection led to a significant reduction in *Yersinia, Chimaeribacter*, and BIGb0383 on days 3 and 7, even in the absence of salt (Figure 3C-D). In parallel, *Leuconostoc* was significantly more abundant in CO_2_-treated fermentations at these timepoints. Similar to the non-treated fermentation, *Weissella* is also increased in low-salt conditions that were treated with CO_2_. Overall, CO_2_ consistently promoted a LAB-dominated, low-Enterobacterales community structure.

These findings demonstrate that CO_2_ injection at the start of fermentation can effectively steer microbial succession, suppress spoilage-associated taxa, and enhance LAB dominance, especially under low- or no-salt conditions. This highlights CO_2_ as a promising tool to modulate fermentation ecosystems toward safer and more desirable outcomes.

## Discussion

This study demonstrates how fermented vegetables can serve as tractable and engineerable model ecosystems for investigating microbial community assembly and succession. By systematically varying environmental parameters, including substrate type, salt concentration, and CO_2_ levels, we observed reproducible patterns of microbial dominance, highlighting how ecological factors can be leveraged to steer microbial communities. These findings reinforce the value of fermented food systems as experimentally accessible platforms for testing ecological hypotheses in complex yet controllable environments.

Understanding bacterial community dynamics and competitive interactions is essential for optimizing the flavor, safety, and quality of fermented vegetables. While previous studies have typically focused on individual vegetable ^12,16,19,47,48^, we performed a standardized longitudinal analysis of the active community across 11 different vegetables. Specifically, the standardized handling was shown to be important as previous work documented that fermentation dynamics were influenced by handling and cutting types of fresh vegetables ^49^.

In our current study, we found that the specific vegetable substrate influenced overall bacterial composition most, while the region of plant (Carposphere, phyllosphere or rhizosphere) also influenced overall bacterial composition, but to a lesser extent. Most fermentations were consistently dominated by LAB, predominantly *Leuconostoc*. Remarkably, tomato and fennel fermentations were dominated by the uncharacterized Enterobacteriaceae taxon BIGB0383, which has also been detected in the *Caenorhabditis elegans* gut ^50^. Given its close phylogenetic relationship to *Pseudescherichia*, an environmental microorganism which functionality remains ambiguous ^51,52^. However, it is important to note that, due to the limitations of the current approach, taxonomic identification was only possible at the genus level. As such, we could not assess the presence of virulence factors or toxins, which would require strain-level resolution.

We then explored the importance of externally controllable factors such as salinity for steering vegetable fermentations. Reducing salt concentration clearly shifted the temporal microbial community dynamics. Lower salt concentrations were associated with higher alpha diversity compared to the reference condition of 2.5% NaCl, accompanied by the presence of less favorable *Enterobacterales* including the genera *Yersinia* and *Chimaeribacter*, as well as the non-*Enterobacterales* genus *Pseudomonas*. A slower microbial succession from heterofermentative *Leuconostoc* to homofermentative *Lactiplantibacillus*-dominated communities was also observed in lower salt concentrations. This trend partially aligned with findings in kimchi, who reported a lower abundance of homofermentative *Latilactobacillus* at lower salt concentrations, although this study was limited to a single replicate^19^. Others have also observed elevated *Enterobacteriaceae* counts at lower salt concentrations in fermented cabbage ^30^, but they only measured colony-forming units of specific taxa of interest while our sequencing approach of the active fraction (RNA) allowed for a more comprehensive overview. Together, our findings clearly demonstrate that reducing salt concentrations not only increases the abundance of unwanted *Enterobacteriaceae* but also prolongs their presence, potentially leading to the accumulation of undesirable compounds such as biogenic amines ^53^.

We also investigated atmospheric conditions by focusing on steering CO_2_ addition as an alternative strategy to modulate this ecosystem. Saturation of the fermentation matrix with CO_2_ in carrot juice successfully induced a more rapid decline of *Enterobacterales* and a clear dominance of *Leuconostoc*. While an immediate pH reduction was observed following CO_2_ introduction, acidification alone could not fully account for the suppression of *Enterobacterales*, as these taxa persisted at much lower pH in control fermentations with 2.5% NaCl. Prior studies have reported the antimicrobial effects of both high-pressure and dissolved CO_2_^32,54–57^, supporting the hypothesis that CO_2_ directly contributed to the observed shifts in community structure. Our results demonstrated that CO_2_ addition consistently decreased alpha diversity, suppressed the growth of potential spoilage organisms, and enhanced the predominance of beneficial *Lactobacillales* across different salt concentrations. Shifts within the LAB community were also observed, including an increase in *Weissella* at lower salt levels. However, a low but non-negligible *Yersinia* concentration was present in various conditions, including the no-salt CO_2_-treated fermentation and the 2.5% salt without CO_2_ injection. This indicates that a combinatorial approach could be beneficial to engineer this ecosystem to minimize the presence of undesirable taxa as early and effectively as possible. Additionally, the addition/flushing of CO_2_ should be validated in various fermentation to reduce possible side effects like the bloater effect in cucumbers ^58^. Overall, CO_2_ addition emerges as a promising strategy to enhance fermentation safety while reducing salt content, opening opportunities for robust and healthier fermented vegetable products.

The results from our current study can be used to explore questions about the ecological mechanisms driving LAB succession in vegetable fermentations. While high salinity has long been considered a key selective pressure favoring LAB, our results suggest that other environmental factors such as the addition of CO_2_ can be used modulate the microbial community assembly. Of note, heterofermentative LAB such as *Leuconostoc* also produce CO_2_ as a metabolic byproduct, which may further also positively influence community dynamics during fermentation.

In summary, this study addressed three key knowledge gaps in the field of vegetable fermentation. First, it showed a systematic LAB dominance of the microbial community across a wide range of vegetable types. Second, it demonstrated the essential role of salt on microbial community dominance and successions. Third, it revealed the potential of CO_2_ addition to simultaneously restore natural microbial dynamics and enable food-safe salt reduction in vegetable fermentations. Given its promising impact, further research into the scalability and robustness of CO_2_ injection in larger fermentation systems is warranted. More broadly than merely food fermentation applications, this study illustrates that targeted manipulation of environmental parameters can be used to steer microbial ecosystems.

## Acknowledgements

The authors would like to acknowledge all colleagues from the Lab of Applied Microbiology and Biotechnology, with special thanks to Nele and Sam for their contributions. We also sincerely thank prof. Paul Cotter for his insightful feedback on this work. Finally, we acknowledge the third-year Bioscience Engineering bachelor students of 2021 and 2022 for their efforts in initiating the vegetable fermentation.

## Data availability

Amplicon sequencing data will be made available on ENA (PRJEB98055) upon publishing.

